# Gaussian accelerated molecular dynamics reveals that a proline-rich signaling peptide frequently samples *cis* conformations when unbound

**DOI:** 10.1101/2021.07.01.450746

**Authors:** Juan Alcantara, Robyn Stix, Katherine Huang, Acadia Connor, Ray East, Valeria Jaramillo-Martinez, Elliott J. Stollar, K. Aurelia Ball

**Author notes:** Correspondence: K. Aurelia Ball.

## Abstract

Disordered proline-rich motifs are common across the proteomes of many species and are often involved in protein-protein interactions. Proline is a unique amino acid due to the covalent bond between the backbone nitrogen and the proline side chain. The resulting five-membered ring allows proline to sample the *cis* state about its peptide bond, which other residues cannot do as readily. Because proline-rich disordered sequences exist as ensembles that likely include structures with the proline peptide bond in *cis*, a robust methodology to accurately account for these conformations in the overall ensemble is crucial. Observing the *cis* conformations of proline in a disordered sequence is challenging both experimentally and computationally. Nitrogen-hydrogen NMR spectroscopy cannot directly observe proline residues, which lack an amide bond, and computational methods struggle to overcome the large kinetic barrier between the *cis* and *trans* states, since isomerization usually occurs on the order of seconds. In the current work, Gaussian accelerated molecular dynamics was used to overcome this free energy barrier and simulate proline isomerization in a tetrapeptide (KPTP) and in the 12-residue proline-rich SH3 binding peptide, ArkA. We found that Gaussian accelerated molecular dynamics, when combined with a lowered peptide bond dihedral angle potential energy barrier (15 kcal/mol), allowed sufficient sampling of the proline *cis* and *trans* states on a microsecond timescale. All ArkA prolines spend a significant fraction of time in *cis*, leading to a more compact ensemble with less polyproline II helix structure than an ArkA ensemble with all peptide bonds in *trans*. The ensemble containing *cis* prolines also matches more closely to *in vitro* circular dichroism data than the all-*trans* ensemble. The ability of the ArkA prolines to isomerize likely affects the peptide’s ability to bind its partner SH3 domain, and should be studied further. This is the first molecular dynamics simulation study of proline isomerization in a biologically relevant proline-rich sequence that we know of, and a similar protocol could be applied to study multi-proline isomerization in other proline-containing proteins to improve conformational diversity and agreement with *in vitro* data.

## 1 Introduction

Proline-rich disordered sequences are one of the common binding motifs for protein-protein interaction domains found in biology. Proline-rich regions are found widely in prokaryotes and eukaryotes and are present in twenty-five percent of human proteins (Kaneko et al., 2008; Williamson, 1994). Despite their prevalence, much is unknown about the structural properties of disordered proline-rich sequences since they are challenging to study both experimentally and computationally. Although a proline-rich region will canonically adopt a polyproline II (PPII) helix when bound to their interaction partner, such as an SH3 domain, these sequences are also flexible and their structures can vary based on the identity of the other amino acids present (Kaneko et al., 2008; Williamson, 1994).

One unique attribute of proline is its ability to isomerize around the peptide bond and sample a *cis* conformation. In the typical *trans* conformation of the peptide bond, the *ω* dihedral angle is 180°, while in the *cis* state, the *ω* angle is 0. The *cis* conformation is energetically disfavored because in this conformation the *α*-carbon atom is positioned on the same side of the backbone as the *α*-carbon for the preceding residue, causing a steric interaction. However, in the case of proline, the unique ring structure means that the *γ*-carbon atom of the proline side chain is bonded to the nitrogen in the backbone, causing an unfavorable steric interaction between the *γ*-carbon and the *α*-carbon of the preceding residue in the *trans* conformation. Thus, the difference in energies between the *cis* and *trans* is reduced for proline when compared to other amino acids, making it thermodynamically possible to find proline residues in the *cis* conformation, while all other residues are very rarely found in *cis*. In the Protein Data Bank (PDB), 5-7% of prolines are found in the *cis* state (MacArthur and Thornton, 1991). A proline residue in a disordered sequence might be expected to sample the *cis* state even more frequently, perhaps in 10% of its equilibrium ensemble, although the identity of the residue preceding the proline will also affect *ω* dihedral angle (Doose et al., 2007; Williamson, 1994). For proline-rich sequences, the typical secondary structure is a left-handed PPII helix, which requires that all peptide bonds in the helix are in the *trans* state. When sequential prolines are in *cis*, a right-handed polyproline I (PPI) helix can form instead (Moradi et al., 2009; William J Wedemeyer et al., 2002). If just a single peptide bond in a disordered sequence is in the *cis* conformation, this results in a kink at that point in the peptide chain.

Although the *cis* and *trans* states for proline are thermodynamically similar, there is a high free energy barrier of 10-20 kcal/mol that must be overcome to switch between the two states (Moradi et al., 2009; William J Wedemeyer et al., 2002). This is primarily due to the partial double bond character of the peptide bond which must be broken to allow rotation around the *ω* dihedral angle. Peptidylprolyl isomerases often catalyze this transition *in vivo* when it is required for folding or protein function (William J Wedemeyer et al., 2002). The uncatalyzed isomerization transition between *cis* and *trans* therefore exhibits slow kinetics, taking place on a time scale of seconds to minutes (Moradi et al., 2009; William J Wedemeyer et al., 2002). For proteins with a stable folded structure, the timescale of the isomerization of a proline residue in the *cis* state can determine the folding rate of the protein (William J Wedemeyer et al., 2002). In some cases, proline isomerization is also important for protein function. For example, the mechanosensing ability of filamin, transcription regulation of the histone H3, and the binding affinity of NCBD to ACTR are all affected by isomerization of a key proline residue (Nelson et al.; Rognoni et al., 2014; Zosel et al., 2018). In the case of proline-rich intrinsically disordered proteins (IDPs), the role of proline isomerization in function has been less well studied. Although many proline-rich disordered sequences have been shown to adopt a conformation with all prolines in *trans* when bound to their interaction partner, these peptides may also interact with their partners when one or multiple proline residues are in *cis*, and how these alternate isomeric states affect binding is largely unknown.

A number of factors contribute to the difficulty of studying the isomerization state of proline in proline-rich disordered proteins. IDP structure is often studied using NMR spectroscopy; however common 2-D protein-NMR experiments using ^1^H and ^15^N are not able to report on proline residues directly because proline lacks an N-H bond that is present in all other amino acids. NOESY experiments and 1-D hydrogen NMR experiments have been performed on polyproline peptides, but it is difficult to specifically identify each residue in such experiments (Best et al., 2007; Kelly et al., 2001). Circular dichroism (CD) reports in an average way on the secondary structure of a protein and the CD ellipticity measurement at 228 nm has been found to be related to the amount of PPII helix present in a peptide (Kelly et al., 2001). However, CD cannot directly report on the *cis* conformations for specific proline residues in a protein’s structural ensemble. FRET and PET can also reveal some information about proline-rich peptide conformations, but only indirectly. Studies using NMR, CD, PET, and FRET have found that polyproline peptide chains sample a significant amount of *cis* conformations in their structural ensembles, which disrupts the canonical PPII structure of these peptides (Best et al., 2007; Doose et al., 2007; Kelly et al., 2001), but little is known experimentally about the *cis* conformational states of more biologically relevant proline-rich disordered sequences.

Computational methods have also encountered challenges in studying the *cis*-*trans* isomerization of the peptide bond in proline-rich proteins. Because of the high energetic barrier between the *cis* and *trans* states, traditional molecular dynamics (MD) simulations cannot capture the isomerization transition, which takes place on a timescale much longer than the longest MD simulations of proteins. Therefore, all traditional MD simulations of IDPs containing proline have kept the isomerization state of the peptide bond constant throughout the simulation, either producing an ensemble where all the prolines are in *trans* or independently simulating a separate ensemble in *cis* and afterward combining the two ensembles with some weight based on the expected time in *cis* (Yedvabny et al., 2014; Zhan and Ytreberg, 2015; Zosel et al., 2018). This method could be fairly effective for a protein containing only one proline residue, although it assumes the probability of this proline being in *cis* is known *a priori*; however, this method is insufficient for many disordered protein interaction sequences that contain multiple proline residues.

In order to account for both *cis* and *trans* isomers of proline in an MD simulation, an advanced sampling method is required to overcome the high energetic barrier between the two states. Researchers have used accelerated MD to overcome the isomerization barrier for very short peptides containing just one proline (Doshi and Hamelberg, 2009; Hamelberg et al., 2005). These studies were important for showing that boosting the potential of low energy states could be an effective way to accelerate the transition between *cis* and *trans* isomers, but did not apply the method to a longer biologically relevant sequence containing multiple proline residues. Other studies have focused on polyproline, a homogenous sequence containing only proline residues, which is not usually the binding motif for protein-protein interaction domains. Using either Monte Carlo sampling or adaptively biased MD and implicit solvent, these studies found that a significant amount of the resulting ensemble contains prolines in *cis* despite the canonical assumption that polyproline sequences adopt a PPII structure with all prolines in *trans* (Moradi et al., 2009; Radhakrishnan et al., 2012; Vila et al., 2004). One Monte Carlo simulation study of a disordered protein ensemble in implicit solvent included *cis* conformations for all proline residues (Martin et al., 2016). Recently, another study has used MD to simulate isomerization in a protein containing multiple proline residues, but for an antibody containing two prolines in a loop region, not a proline-rich disordered sequence (Masiero et al., 2020). These results suggest that important proline-rich signaling peptides also likely sample ensembles containing *cis* conformers, and accounting for these structures in MD simulations could be important for understanding proline-rich IDP function.

We focus on the proline-rich peptide ArkA, a disordered region of the yeast actin patch kinase Ark1p. This proline-rich IDP helps regulate the actin cytoskeleton assembly by binding to an SH3 domain of Actin Binding Protein 1 (Abp1p). The key binding region of Ark1p is 12 residues in length (K_(3)_P_(2)_T_(1)_P_(0)_P_(−1)_P_(−2)_K_(−3)_P_(−4)_S_(−5)_H_(−6)_L_(−7)_K_(−8)_) and contains five prolines. In an NMR structure of ArkA bound to the SH3 domain, all five prolines are in the *trans* conformation and the N-terminal proline-rich region of the peptide adopts a PPII helix (Stollar et al., 2009). However, when this sequence is not bound to the SH3 domain, the proline residues may indeed sample *cis* conformations in equilibrium with the all-*trans* state. Recently we have published a paper on the binding of ArkA to the Abp1p SH3 domain, but in those simulations, the large energy barrier for isomerization of the peptide bond prevented us from sampling any conformations containing *cis* prolines (Gerlach et al., 2020). Here, we attempt to capture the complete conformational ensemble of unbound ArkA by using Gaussian accelerated MD (GaMD) to overcome this barrier and simulate numerous transitions between the *cis* and *trans* states for each proline (Miao et al., 2015). We find that in order to achieve adequate sampling of the transition we must first lower the potential barrier between *cis* and *trans* for the peptide bond before running GaMD simulations. The resulting ArkA ensemble contains a significant population of structures with one or more proline residues in *cis*, which breaks up the PPII helix structure. This GaMD generated ensemble is also more consistent with *in vitro* CD data than an ensemble containing only the *trans* isomer. In future studies of proline-rich disordered sequences like ArkA, it is important to include *cis* proline conformational states in the unbound ensemble and consider what role these conformations have on binding to partner proteins and biological function.

## 2 Materials and Methods

### 2.1 Structures and simulations overview

GaMD simulations (Miao et al., 2015) were conducted on three different peptide systems: a KPTP tetrapeptide with the Amber ff14SB force field (default barrier), KPTP with a modified Amber ff14SB force field that has a lowered potential barrier between the *cis* and *trans* conformers of the peptide bond in the dihedral angle potential energy term (lowered barrier), and a 12-residue segment of ArkA with the lowered barrier (Scheme 1). The residue numbering for ArkA is based on the standard system (Lim et al., 1994). The ArkA peptide simulations were run with a fully extended peptide starting structure, or with a starting structure from the NMR structure of ArkA bound to the Abp1p SH3 domain (Stollar et al., 2009) (PDB: 2RPN). Both starting structures were capped with an acetyl group at the N-terminus and amino group at the C-terminus to neutralize terminal charges. The KPTP simulations were conducted before the ArkA simulations to determine which potential energy function would be used for the peptide bond dihedral in the ArkA simulations to increase the likelihood of capturing proline isomerization. Overall simulation lengths and relevant properties are listed in Table 1.

**Scheme 1.**
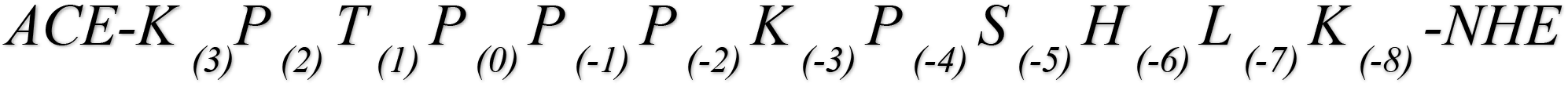
ArkA peptide sequence and numbering.

**Table 1.**
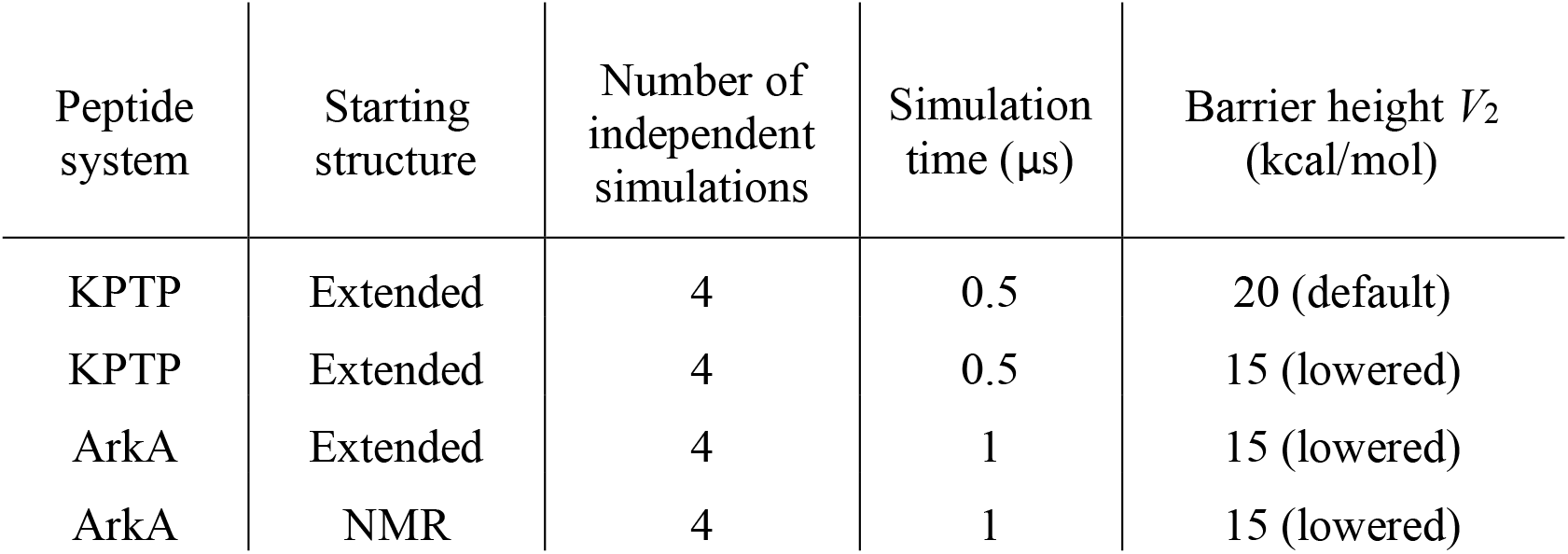
Summary of simulations run. Barrier height denotes the parameter *V_2_* used in the peptide bond dihedral function that determines the potential energy barrier between the *cis* and *trans* states.

| Peptide system | Starting structure | Number of independent simulations | Simulation time (μs) | Barrier height $V_2$ (kcal/mol) |
| --- | --- | --- | --- | --- |
| KPTP | Extended | 4 | 0.5 | 20 (default) |
| KPTP | Extended | 4 | 0.5 | 15 (lowered) |
| ArkA | Extended | 4 | 1 | 15 (lowered) |
| ArkA | NMR | 4 | 1 | 15 (lowered) |

### 2.2 GaMD simulations

Simulations were conducted on a Linux cluster using Amber 18 pmemd CUDA for GPU functionality (Case et al., 2018). GaMD was used for enhanced sampling and to overcome the barrier between the *cis* and *trans* states of the *ω* angle (Miao et al., 2015). The Amber *LEaP* module was used to create topology and initial coordinate files for all simulated starting structures. The simulations were run with the Amber ff14SB force field (Maier et al., 2015). For the lowered barrier simulations, a modification was made to the *V_2_* constant in the peptide bond dihedral potential function. This adjustment was made similarly to the way that Doshi and Hamelberg increased the *V_2_* constant to increase the barrier between the *cis* and *trans* states of proline (Doshi and Hamelberg, 2009). The *V_2_* constant in these force fields is solely responsible for determining the energy barrier height for peptide bond (*ω* angle) isomerization and does not affect other dihedral angles or any other conformational sampling coordinate. *V_2_* was decreased from the default of 20 kcal/mol to 15 kcal/mol. Simulations were solvated with the TIP3P-FB water model (Wang et al., 2014), such that the edge of the box was at least 15 Å from any atom in the peptide. Chloride ions were added to neutralize the system (1 for KPTP and 3 for ArkA).

Systems were minimized in two rounds with the first one placing a harmonic potential restraint on the peptide of 10 kcal/mol/Å^2^. Systems were then heated from 100 K to 300 K using a 2 fs integration time step for 20,000 steps (40 ps). Equilibration was also performed in two rounds: the first with a 10 kcal/mol/Å^2^ restraint and pressure and density convergence to 1.013 bar with the Berendsen barostat for 50 ps; the second was run without the restraint for 500 ps and the barostat was switched to a Monte Carlo barostat to maintain isobaric conditions. A final equilibration was performed for 52 ns to calculate the GaMD potentials that dictate how the potential energy landscape will be boosted for enhanced conformational sampling during the production stage. Boosts were calculated and applied with respect to the total potential of the system (igamd = 3) with an energy threshold boost biasing value equal to the upper bound of the potential energy (iE =2). A 9 kcal/mol upper limit was set for the total potential acceleration standard deviation for the purposes of reweighting the produced ensembles (σ_0P_). Productions were conducted under isobaric conditions with a 2 fs time step and a 0.1 ps frame output in order to ensure the ensembles could be appropriately reweighted once completed. Independent simulations were started with new random velocities. Bonds to hydrogen were constrained using the SHAKE algorithm during all simulations. The particle-mesh Ewald procedure was used to handle long-range electrostatic interactions with a non-bonded cutoff of 9 Å for the direct space sum.

The NMR structure GaMD equilibration step experienced a random jump in potential energy (Figure S1), slightly increasing the applied GaMD energy boost relative to the extended structure simulations (a known problem with the Amber 18 implementation of GaMD). Because of this, the extended structure production steps were rerun with the boost potentials calculated in the NMR GaMD equilibration to maintain consistency between the two sets of simulations and improve sampling of the peptide bond isomerization. The random boost was not substantial enough to create a physically irrelevant potential energy landscape, so any abnormally high energy structure sampled was adjusted for during the reweighting process.

### 2.3 Trajectory analysis

The *cpptraj* module in AmberTools 18 was used for all trajectory analysis (Case et al., 2018). In house Python scripts were used for all data processing and plots. Visual Molecular Dynamics was used to create structural figures (Humphrey et al., 1996).

Simulations were reweighted to create a potential of mean force (PMF) for the proline *ω* dihedral angles using both the cumulant expansion on the second order and Maclaurin series expansion to the 10th order provided by the PyReweighting tool kit (Miao et al., 2014). The cumulant expansion on the second order more accurately reproduced the free energy barriers to isomerization, but the Maclaurin expansion method allowed us to reweight independent of the reaction coordinate. Therefore, all other equilibrium data presented from the GaMD simulations have been reweighted using the Maclaurin expansion series to the 10^th^ order. The PyReweighting tool kit was also used to calculate the anharmonicity along the proline *ω* dihedral for the GaMD simulations. An average *ω* angle anharmonicity value < 10^−3^ kcal/mol was used as the cutoff to determine whether adequate sampling had been reached (Miao et al., 2015).

Histograms for the *ω* angles in the KPTP peptide simulations were created to visualize the distributions of angles sampled during the simulations to define ranges for *cis* and *trans* values for the subsequent ArkA analysis. Two *ω* angle states close to the canonical 0° and 180° *cis* and *trans* values were observed in the KPTP system (Figure S2). Based on the distribution of the *ω* angles in KPTP, the ranges *ω* = −90° to +50° and *ω* = +100° to +240° were used to define the *cis* and *trans* states, respectively. Percentages of time each proline spent either in *cis* or *trans* as well as the flipping frequencies between them were calculated for KPTP and ArkA simulations.

Proline correlation analysis was performed to compare the likelihood of multiple prolines being physically dependent on one another to isomerize to either cis or trans. All possible combinations of prolines were analyzed for correlation in the KPTP and ArkA simulations. Expected percentages of time each group of prolines sampled *cis* together were calculated by assuming such events were independent of each other and thus equal to the product of their individual *cis* sampling times. These expected values were then compared to the actual percentages of time the proline combinations sampled *cis* simultaneously within the simulations.

Bend, turn, and 3_10_ helix secondary structures for ArkA were calculated using the DSSP algorithm in *cpptraj*. PPI and PPII helix structures were calculated based on canonical values of *ϕ*, *ψ*, and *ω* angle ranges as shown in Table 2 (Mansiaux et al., 2011; Radhakrishnan et al., 2012). Running averages of bend, turn, 310 helix, and end-to-end distance were used to test structure convergence between extended and NMR starting structure systems of ArkA. Every 10th frame (every 1 ps) of the ArkA trajectories was used for testing convergence. Proline ring pucker states were calculated using the *χ^2^* dihedral angle (C*α*—C*β*—C*γ*—C*δ*) and defined by canonical ranges shown in Table 3 (Radhakrishnan et al., 2012).

**Table 2.** Canonical backbone dihedral angles that define PPI and PPII. Note that the *ω* angle range for PPII could also be written as +100° to +240°.

| Dihedral angle | PPI (°) | PPII (°) |
| --- | --- | --- |
| $\phi$ | -104.6 to -46.6 | -104.6 to -46.6 |
| $\psi$ | +131 to +189 | +107.9 to +165.9 |
|  | -90 to +50 | +100 to +180<br>or |
| $\omega$ | | -180 to -120 |

**Table 3.** Proline side chain dihedral angle *χ*_2_ defines the canonical proline ring pucker states.

| Pucker Type | $\chi_2$ (°) |
| --- | --- |
| Up pucker | > +10 |
| Down pucker | < -10 |
| Planar | -10 to +10 |

### 2.4 Replica exchange data

Replica Exchange MD (REMD) ArkA simulations previously run as a starting point for binding simulations were used to construct a comparison structural ensemble that contains only *trans* proline conformations (Gerlach et al., 2020). REMD simulation lengths and structures are listed in Table 4. Structures were saved every 25 ps and the first 50 ns of each simulation was not used for analysis. Data from NMR and extended structure starting systems were combined and analyzed together.

**Table 4.** 12-residue ArkA REMD simulation data adapted from Gerlach *et al*. 2020.

| Peptide system | Starting structure | Number of independent simulations | Total simulation time amongst independent simulations (μs) |
| --- | --- | --- | --- |
| ArkA | NMR | 3 | ~0.5 |
| ArkA | Extended | 3 | ~0.5 |

### 2.5 Computational calculation of CD spectra

CD can be used to differentiate between different secondary structure compositions of a protein. The SESCA CD program (v0.95) was used to calculate theoretical CD spectra for the ArkA simulation ensembles (Nagy et al., 2019). The DS5-4SC1 basis-set was chosen to calculate the theoretical spectra for ArkA, primarily because of its Turn 1 secondary structure definition containing a PPII component, which allows the calculated spectrum to differentiate between ensembles with different amounts of PPII content.

An ensemble containing every 100th frame (10 ps) of trajectory data from the ArkA GaMD simulations was used for the SESCA CD spectra calculations. The REMD simulations were also analyzed with SESCA. Percentages of basis-set secondary structure compositions were also calculated. A scaling factor for the *in vitro* ArkA CD spectrum was calculated from the REMD ensemble to correct for measurement cell concentration errors that may have influenced the measured CD spectrum. However, to compare all theoretical spectra to the experimental one, and to avoid modifying the experimentally measured values, the inverse of this scaling factor (0.510) was applied to all theoretically calculated CD spectra.

PPII content percentages were also extracted from the CD data using an approximation that calculates the composition of PPII in a peptide,

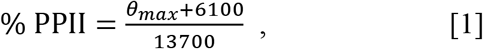

based on the CD ellipticity (10^3^ deg cm2/dmol) at the 228 nm wavelength (*θ_max_*) (Kelly et al., 2001). Average PPII percentages from the *in vitro* and calculated CD spectra were compared to average PPII percentages calculated directly from the simulation trajectory data.

### 2.6 CD spectroscopy of ArkA peptide

The 12-residue ArkA peptide (KPTPPPKPSHLK) was synthesized by Peptide 2.0 with acetylated N-terminus and amidated C-terminus and purified to >95% purity. The peptide was resuspended and dialyzed against 5 mM Sodium Phosphate buffer, pH 7.0. CD spectra were collected using a Chirascan™-plus CD spectrometer with a 250 μL 70 μM sample at a pathlength of 1 mm. Measurements were obtained at 2 seconds/point for wavelengths between 180 nm – 260 nm with a step size of 1 nm. All spectra were blank corrected with buffer only (5 mM Sodium Phosphate buffer, pH 7.0). Data were collected in millidegrees and were converted to degrees and mean residue ellipticity, [*θ*]_*MR*_, according to

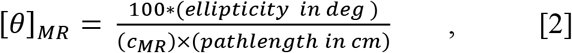

where the mean residue molar concentration, *C_MR_*, is calculated based on the molar concentration, *c*,

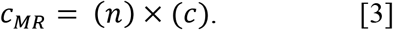

The number of residues, *n*, equals 12 in the case of ArkA.

## 3 Results and Discussion

In order to understand the role that peptide bond isomerization has in a proline-rich IDP’s structural ensemble, a method that can overcome the large energy barrier between the *cis* and *trans* states to adequately sample isomerization must be implemented. We chose to use the GaMD method because it does not require defining a reaction coordinate, can be implemented in explicit solvent, and allows for accurate reweighting of the resulting conformations to recover the unbiased ensemble (Miao et al., 2015). To determine whether GaMD is capable of sampling proline isomerization in the SH3 binding peptide ArkA, the short tetramer peptide KPTP (consisting of the first four residues of ArkA) was simulated as a test before simulating the longer ArkA (12-residues).

### 3.1 Proline isomerization of tetramer KPTP using GaMD

Initially, GaMD simulations were run on KPTP using the Amber ff14SB force field (Maier et al., 2015), which contains a peptide bond torsional potential term with a 20 kcal/mol barrier between the *cis* and *trans* minima. This potential barrier has been shown to be too low to reproduce the experimentally observed free energy barrier between the *cis* and *trans* states (Doshi and Hamelberg, 2009). However, it is still a large barrier to overcome in MD simulations, and even using the GaMD method we only saw isomerization occur on a time scale of around once every 8 ns (Figure 1A). Therefore, to increase sampling and because we were much more interested in the conformations in the *cis* and *trans* energy minima than in the energies during the transition, we implemented a lowered potential barrier for peptide bond isomerization of 15 kcal/mol (see GaMD simulations in methods). GaMD simulations of KPTP with the lowered barrier resulted in an increase of proline isomerization frequency of more than an order of magnitude (Figure 1B). Both the default barrier and lowered barrier simulations showed that the proline peptide bond *ω* dihedral angle can clearly occupy two distinct states that are similar to the canonical *cis* and *trans* values (Figure S2).

**Figure 1.**
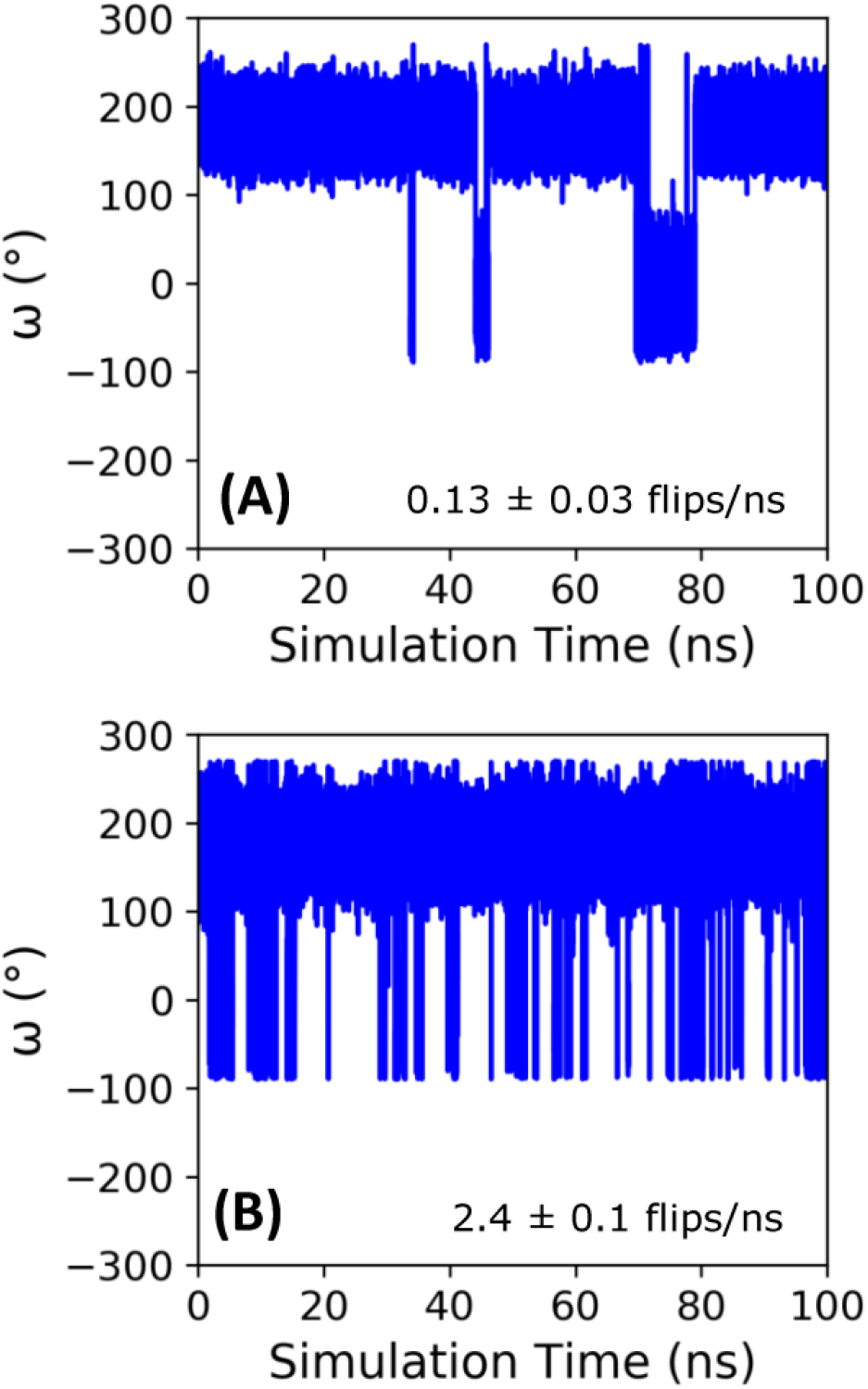
Time course of the *ω* angle for the first proline in KPTP from 100 ns of a single GaMD simulation using the default **(A)**, and lowered barrier force fields **(B)**. The *ω* angle is around 180° when in the *trans* state and around 0° when in the *cis* state. The isomerization frequency represents the average over all independent simulations.

Cumulant expansion on the second order was used to reweight the GaMD ensembles and construct a PMF along the *ω* angle reaction coordinate (Figure 2A). Very high free energy values at the barriers between the minima for the default barrier simulations indicate that sampling of the transition is insufficient to accurately assess the barrier height. However, with the increased isomerization in the lowered barrier simulations, the barrier height falls closer to where we expect, between 10 and 15 kcal/mol, and the *cis* and *trans* minima for both sets of simulations remained relatively equivalent in energy. Average anharmonicity (Anharm_Def_ = −0.0013, Anharm_Low_ = −0.0014) values for both systems were below the cutoff of 10^−3^, indicating that accurate reweighting of the boosted ensemble is possible using cumulant expansion on the second order (Figure S3). We also used Maclaurin series expansion to the 10th order for reweighting, which results in a smoother PMF plot, but tends to underestimate free energy barrier heights and can also result in slight shifts in the location of minima (Figure 2B). While the free energy barriers in this PMF are underestimated, the depth and location of the *cis* and *trans* minima are similar to the PMF found using cumulant expansion on the second order. Maclaurin series expansion reweighting also has the advantage that the resulting weights are reaction coordinate independent and can be applied to all further analyses of the ensemble. Therefore, we used the Maclaurin series expansion reweighting for all subsequent analyses of both the KPTP and ArkA conformational ensembles. Although the isomerization rates were different for the default and lowered barrier simulations, the percent of the ensemble in *cis* for each proline (after reweighting) was similar in both sets of simulations (first proline: ~14% for both, second proline: ~10% for both) (Table S1).

**Figure 2.**
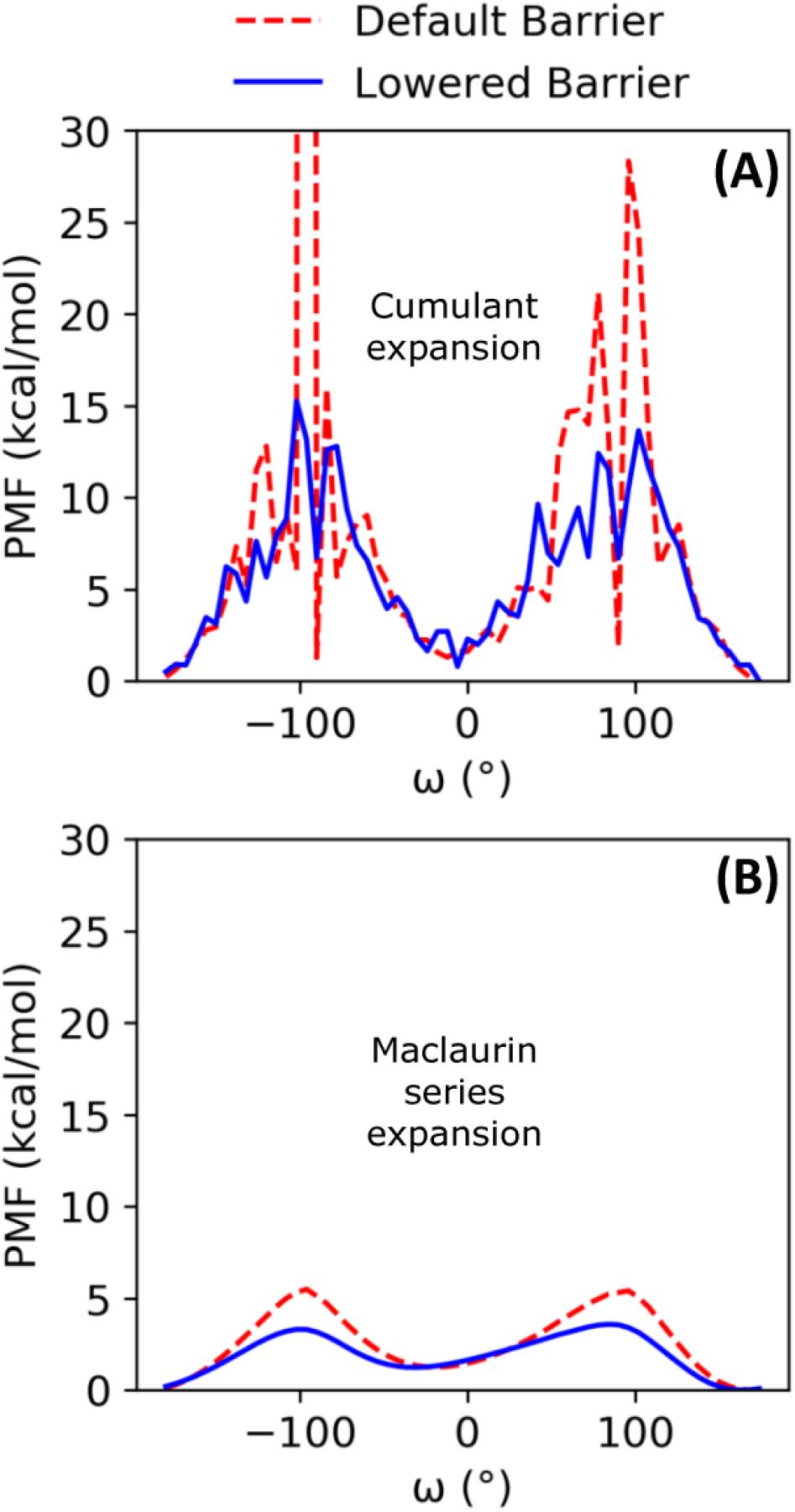
PMF free energy landscapes over the *ω* dihedral angle for GaMD default and lowered barrier KPTP simulations for the first proline reweighted using **(A)** cumulant expansion on the second order and **(B)** Maclaurin series expansion.

### 3.2 Proline isomerization of the proline-rich IDP ArkA

To simulate proline isomerization for the 12-residue ArkA peptide, we chose to use the lowered potential energy barrier to represent the peptide bond dihedral angle since it allowed more frequent isomerization with GaMD and therefore improved sampling of the isomer states. Simulations were conducted using two different starting structures, an extended structure and an NMR structure taken from the AbpSH3 bound state (PDB: 2RPN) (Stollar et al., 2009), to assess sampling by comparing the resulting ensembles. Both starting structures contained all peptide bonds in the *trans* state. Recently published work using the same extended and NMR starting structures, but using the REMD enhanced sampling method, did not capture any *cis* sampling (Gerlach et al., 2020), which is consistent with other studies that show REMD is not an effective method for observing proline isomerization (Neale et al., 2016; Yedvabny et al., 2014; Zosel et al., 2018).

In the GaMD simulations, all five ArkA prolines isomerized with a high frequency, sampling both the *cis* and *trans* state (Table S2). Turn and 310 helix secondary structure, as well as end-to-end distance, showed that the two sets of simulations started to converge over time, though additional simulation time would be needed for complete sampling (Figure S4). Because the extended and NMR starting structure simulations were not completely converged, the extended structure simulations have slightly higher *cis* content for each proline (Figure 3). However, the *cis* percentage differences were small enough that we combined the data from all ArkA simulations into one ensemble for further analysis. The PMF and anharmonicity values for both GaMD extended and NMR simulations show sampling was adequate for accurate reweighting of the ensemble (Figure S5). These results show that using GaMD with a lowered peptide bond dihedral angle potential energy barrier allows for adequate sampling of the proline *cis* state for all prolines in ArkA on the nanosecond to microsecond time scale and that this method would likely work for other biologically relevant proline-rich peptides. In contrast to a previous Monte Carlo simulation of the 81-residue disordered transcription factor Ash1 in implicit solvent (Martin et al., 2016), several of the ArkA prolines are in *cis* in more than 10% of the ensemble. The higher *trans* occupancy for the Ash1 prolines might be due to the larger size of the disordered region, differences in adjacent residues to the prolines, or differences in the simulation method.

**Figure 3.**
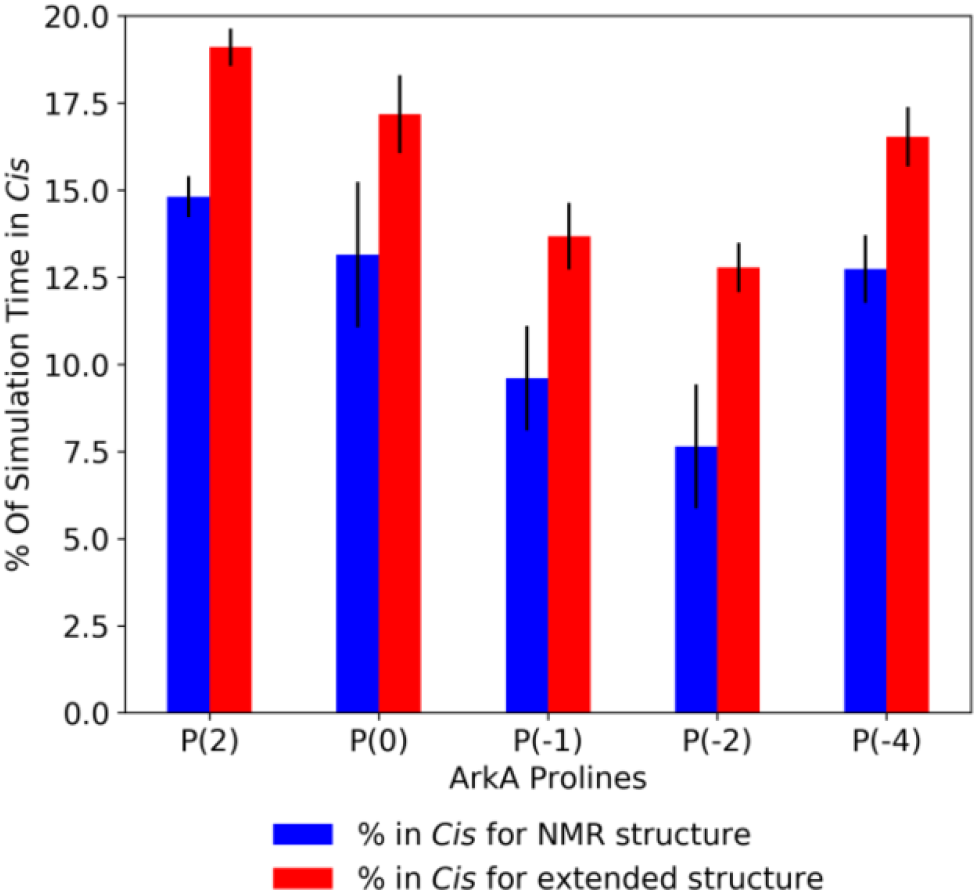
Average ArkA proline *cis* state percentages in the GaMD simulations, for the NMR (left bar, blue) and extended (right bar, red) starting structures. Error bars represent the standard deviation over all independent simulations for the given starting structure.

In order to visualize the ArkA conformational ensemble, we plotted the probability density based on the number of prolines in *cis* and the peptide end-to-end distance (Figure 4). Using these reaction coordinates, the most populated ArkA conformational state has an end-to-end distance of ~24 Å with no *cis* prolines (Figure 4). However, the ensemble also contains many conformations with multiple prolines in the *cis* conformation simultaneously. In fact, with five proline residues that can sample both *cis* and *trans* isomers, ArkA only spends 36% (±7%) of the GaMD ensemble with all peptide bonds in trans. There is also an inverse relationship between the number of prolines in *cis* and how extended the peptide structure is, since the *cis* conformation introduces a kink into the peptide chain, as has been previously observed for polyproline peptides (Moradi et al., 2009; Radhakrishnan et al., 2012). These kinks and slight end-to-end shortening can be observed in the example peptide snapshots (Figure 4).

**Figure 4.**
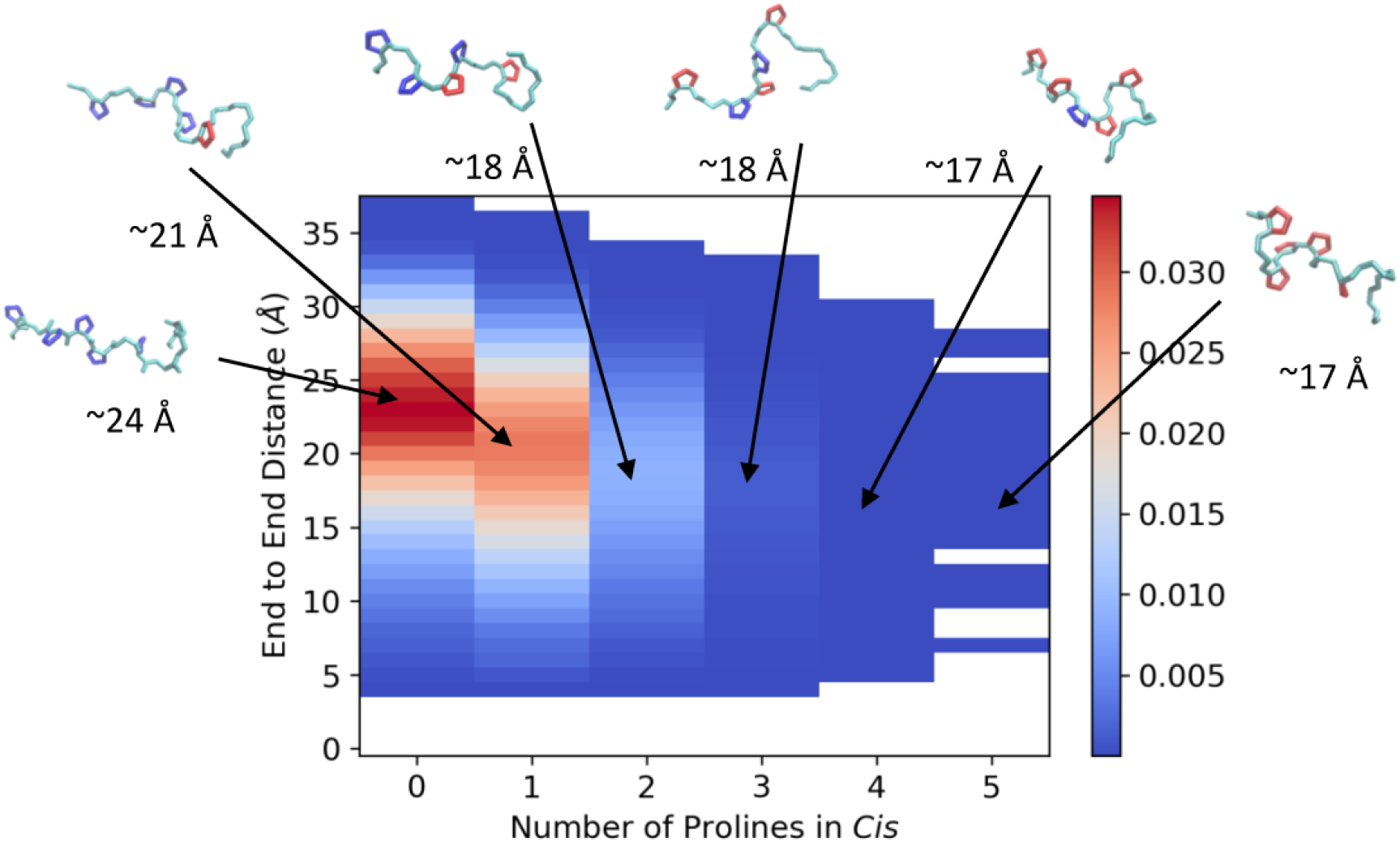
Joint distribution showing the probability of conformations based on end-to-end distance and number of proline residues in *cis* for the combined ensemble from the GaMD extended and NMR structure simulations of ArkA. Example peptide snapshots are oriented such that the N-terminus is pointed leftward. The backbone is depicted in cyan, *trans* prolines in blue and *cis* prolines in red. Approximate end-to-end distances for the representative snapshots are included below each.

The peptide bond dihedral angle of proline affects the overall structure of the ArkA peptide. In the ensemble generated using GaMD, ArkA samples more bend and PPI helix structure than in the previously published REMD ensemble (Gerlach et al., 2020), while spending less time in PPII and 310 helix (Figure 5). PPI helix was not observed in the REMD systems because it requires a *cis ω* angle (Figure 5D), but is a stable structure for polyproline sequences when all the prolines are in *cis* and has been observed in simulations of polyproline (Moradi et al., 2009; William J Wedemeyer et al., 2002). The increase in the bend character of ArkA in the GaMD simulations compared to the REMD ensemble is also likely due to the kinks in the peptide structure caused by the presence of *cis* prolines. Looking at the dihedral angles of the proline side chain, we also see that the down-pucker state of proline is more prevalent than the up-pucker state in the GaMD ensemble (Table S3), consistent with previous studies that show *cis* isomers favor this pucker state while *trans* isomers have no preference (Radhakrishnan et al., 2012).

**Figure 5.**
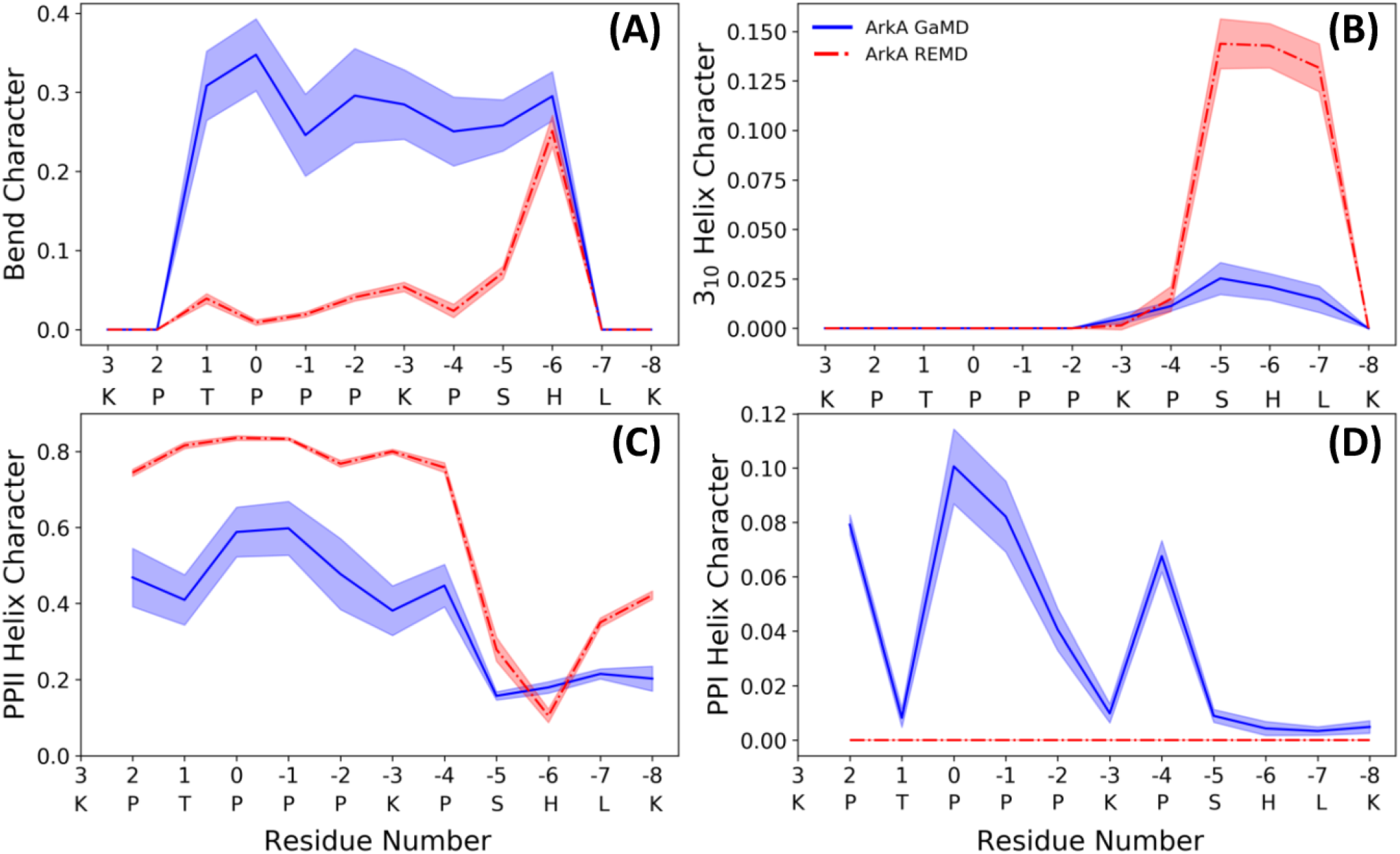
ArkA secondary structure composition for **(A)** bend, **(B)** 310 helix, **(C)** PPII helix, and **(D)** PPI helix. The fraction of the ensemble in the structure is plotted by residue for both the GaMD ensemble (blue solid line), which contains *cis* proline conformations, and the REMD ensemble (red dashed line), which contains only *trans* prolines. The shaded errors represent the standard deviation over all independent simulations.

Because ArkA contains multiple prolines that can isomerize, we also wanted to test whether there is any correlation between their isomeric states, such as two adjacent prolines being more likely to sample *cis* at the same time. To do this, we calculated the expected joint probabilities of each pair of prolines being in *cis* simultaneously assuming that every proline’s isomeric state were independent of the other residues. These calculated probabilities were then compared to the observed simultaneous *cis* occurrences (Figure 6), but no significant difference was found. We also looked for correlations among more than two prolines, but still no significant correlations were detected (Figure S7). This indicates that there is no strong cooperative effect based on proline ring stacking or other interactions that make it more favorable for consecutive prolines to adopt the same structure, and instead, the isomerization is purely stochastic. A similar result has been observed for polyproline sequences in Monte Carlo simulations (Radhakrishnan et al., 2012; Vila et al., 2004). It is possible that other proline-rich peptides might exhibit more correlations between the proline residues depending on the adjacent sequence. It has been shown in several studies that the identity of surrounding amino acids will affect the relative stability of the proline *cis* and *trans* isomers (Brown and Zondlo, 2012; Hamelberg et al., 2005; MacArthur and Thornton, 1991). Additionally, more distal sequence effects could be present in the context of the full-length protein. Regardless, the combinatorial effect of uncorrelated proline isomerization suggests that biologically relevant proline-rich disordered sequences will, in general, contain a significant number of structures with *cis* isomers based on the combined independent probabilities of *cis* for each proline residue.

**Figure 6.**
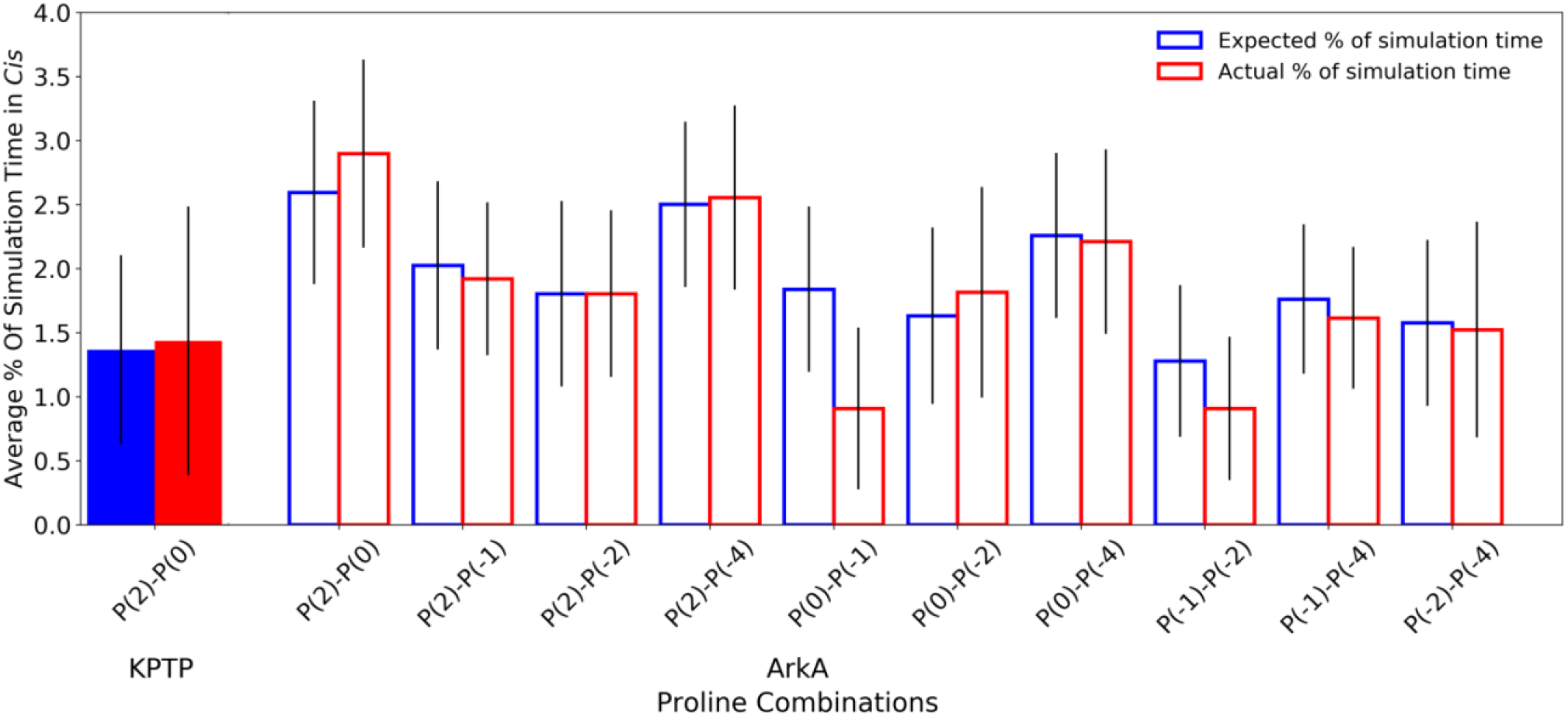
Pair-wise proline *cis* state correlations in the KPTP (solid bars) and in the ArkA GaMD (open bars) systems. Error bars represent the standard deviation between independent simulations. Calculated expected percentages (on the left in blue) are calculated based on the individual *cis* percentages for each proline assuming all proline pairs sample *cis* independently of each other.

### 3.3 Effect of *cis* conformations on ArkA CD spectrum

CD was used to compare the ArkA secondary structures sampled in our simulated ensembles to those observed *in vitro*. Although CD can be used to determine the amount of α-helix and β-sheet in a protein, it is difficult to calculate the exact amount of specific secondary structure content in an IDP ensemble based on CD data because there are multiple possible structural ensembles that would be consistent with the same CD spectrum, especially when more secondary structure types besides α-helix and β-sheet are considered (Greenfield, 2007; Johnson, 1999; Louis-Jeune et al., 2012; Micsonai et al., 2015; Sreerama et al., 2000). Therefore, we chose to calculate theoretical CD spectra based on our simulated ensembles using the SESCA program (Nagy et al., 2019). These theoretical spectra can then be compared to *in vitro* data. The SESCA DS5-4SC1 basis set was chosen for the calculation because it includes a secondary structure category, Turn 1, that includes PPII helix turns, and therefore can be used to differentiate between structures with and without *cis* prolines.

We calculated theoretical CD spectra using both our ArkA GaMD ensemble and the all-*trans* REMD ensemble and compared both to *in vitro* data on the peptide (Figure 7). The calculated spectrum from the GaMD ensemble shows a closer resemblance to the *in vitro* data in both the wavelength and ellipticity intensities compared to the REMD ensemble spectrum, suggesting that *cis* proline isomers are present in the *in vitro* ArkA conformational ensemble. To understand why the spectrum calculated from our GaMD ensemble is closer than the REMD ensemble to the *in vitro* CD spectrum, we examined the secondary structure percentages that were used to generate the calculated spectra. For both the GaMD and REMD ensembles, “Turn 1” is the most common secondary structure (Table 5). The average Turn 1 character for the REMD ensemble is much larger than the GaMD ensemble since it contains more PPII helix with all of the prolines in *trans*. Instead, the GaMD ensemble contains more “Turn 2” and “Other” structure, which may include PPI helix and other conformations with one or more prolines in *cis*.

**Figure 7.**
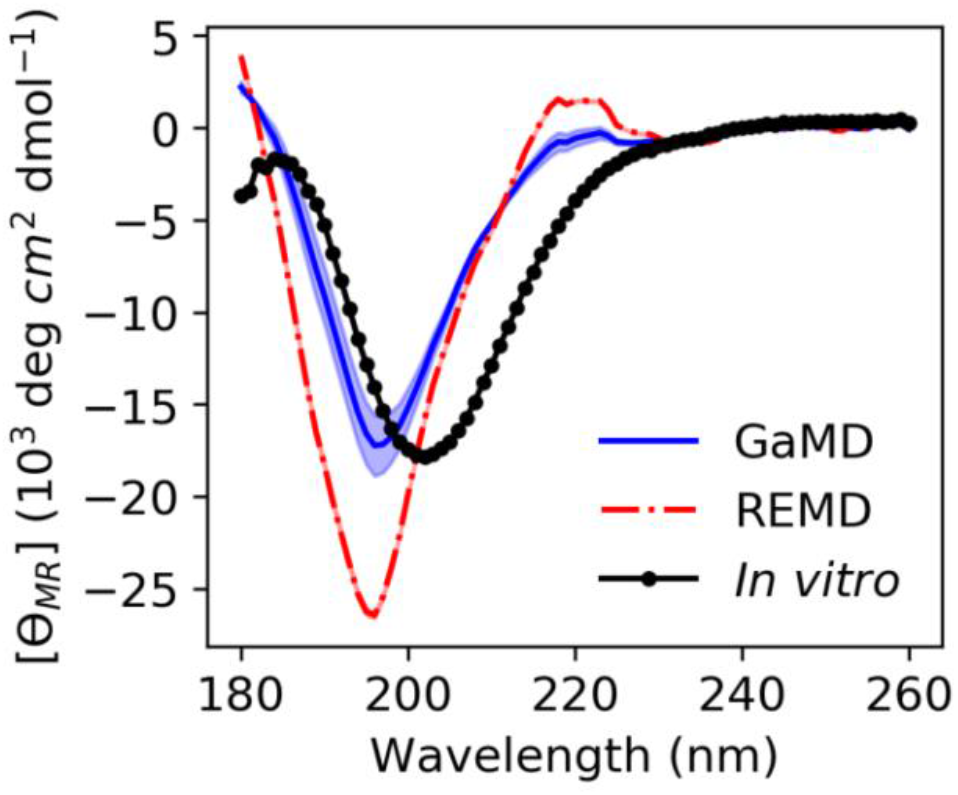
ArkA *in vitro* CD spectrum (black solid line with data points as circles) compared to the calculated theoretical CD spectrum based on the ArkA GaMD (solid blue line) and REMD (dashed red line) ensembles. The theoretical spectra were calculated using SESCA (Nagy et al., 2019) and scaled by a factor of 0.51. The shaded region represents the standard deviation over all independent simulations for each system (the REMD shaded region is the same width as the line).

**Table 5.** Average CD secondary structure compositions for GaMD and REMD systems as defined by the DS5-4SC1 basis-set.

| DS5-4SC1 basis set structures | GaMD simulation percentage | REMD simulation percentage |
| --- | --- | --- |
| Turn 1 | $45 \pm 7$ | $82.7 \pm 0.3$ |
| Turn 2 | $26.5 \pm 0.8$ | $12.4 \pm 0.3$ |
| Other | $25 \pm 6$ | $4.7 \pm 0.2$ |
| Helix 1 | $1.3 \pm 0.5$ | $0.10 \pm 0.08$ |
| Beta 1 | $1.1 \pm 0.3$ | $0 \pm 0$ |
| Helix 2 | $0.60 \pm 0.04$ | $0.12 \pm 0.07$ |

Ellipticity intensity minimums for the GaMD and REMD ensemble are shifted to a lower wavelength (196 nm) compared to the *in vitro* ensemble (202 nm). This can be explained by examining the basis set spectra used to construct the theoretical ensembles. According to the basis spectra for DS5-4SC1 in Figure S12 of (Nagy et al., 2019), the Turn 1 spectrum has a minimum at 195 nm and maximum at 218 nm, while the Turn 2 spectrum has a minimum at 202 nm and no maximum at 218 nm. The calculated GaMD ensemble spectrum has a smaller minimum at 196 nm and smaller maximum at 218 nm than the REMD ensemble spectrum because it has less Turn 1 content; however, it may still contain more Turn 1 content than the *in vitro* spectrum. Since Turn 1 includes PPII conformations, this indicates that our calculated spectra for the GaMD ensemble may still contain more PPII conformations and less *cis* proline conformations than the true *in vitro* ArkA ensemble. Additionally, the SESCA basis sets were all developed for proteins with peptide bonds in *trans* and a basis set that explicitly takes into account *cis* prolines and PPI structure might be able to more clearly identify differences between the *in vitro* and MD ensembles.

We also used another method to directly calculate the amount of PPII structure in the ensemble from the CD data developed by Kelly and coworkers (Kelly et al., 2001). This empirical method based on data from model proteins with varying degrees of PPII character uses CD ellipticity values at wavelengths of 228 nm to calculate the PPII percentage of the sample (Equation 1) (Kelly et al., 2001). This equation was used to obtain the PPII percentage from the *in vitro* CD spectrum and the calculated CD spectra from our GaMD and REMD ensembles. For the MD ensembles, this number was compared to the actual PPII percentage calculated directly from the simulated structures. Based on the ellipticity at 228 nm, the PPII percentage from the GaMD ensemble is relatively consistent with the PPII composition calculated from both the GaMD and the *in vitro* CD spectra, while the REMD ensemble PPII composition is considerably higher (Table 6). Overall, our CD analysis suggests that an ArkA ensemble that contains a significant population of *cis* proline conformations (as calculated by GaMD) more closely resembles the true *in vitro* ArkA ensemble.

**Table 6.** Percent averages for PPII in the GaMD and REMD systems obtained from averaging per-residue PPII ratios over the whole peptide (Figure 5), and calculated from CD data using Equation 1. The average ellipticity values at 228 nm used to calculate PPII% are also included.

| Simulation Systems | PPII % from structure data | PPII % from CD ellipticity at 228 nm | Ellipticity at 228 nm ($10^3 \text{ deg cm}^2\text{dmol}^{-1}$) |
| --- | --- | --- | --- |
| GaMD | $29 \pm 4$ | $38.7 \pm 0.6$ | $\sim -0.796$ |
| REMD | $55.9 \pm 0.3$ | $42.1 \pm 0.1$ | $\sim -0.334$ |
| *In vitro* Ensemble | - | 35.5 | -1.30 |

## 4 Conclusion

Using the SH3 domain binding IDP ArkA as a model peptide, we have performed the first, to our knowledge, MD simulations of a biologically relevant proline-rich sequence that exhibits isomerization of multiple proline residues. We are able to overcome the large free energy barrier between the *cis* and *trans* states of proline by using the GaMD advanced sampling technique as well as a perturbed peptide bond dihedral angle potential with a lower energy barrier. Using this method, we observe many isomerization events on the microsecond time scale of our simulations allowing us to thoroughly sample all possible combinations of proline residues in *cis* and *trans*. For the ArkA system, the computational time to perform a one-microsecond GaMD simulation on a GPU was a few days, allowing for good sampling of the ArkA ensemble on a practical time scale. This method could likely be implemented for other disordered proline-rich sequences in the future to achieve a more realistic picture of their conformational ensembles that includes the *cis* conformation of the peptide bond dihedral.

After using GaMD to sample the ArkA conformational ensemble, we find that this ensemble contains a large number of structures with one or more of the five ArkA proline residues in *cis*, and only about a third of the ensemble consists of all-*trans* conformations. The inclusion of these *cis* proline conformations also provides a better match between our MD ensemble and *in vitro* CD data collected on ArkA. The *cis* peptide bond angle introduces a kink into the ArkA structure, breaking up the PPII helix in the N-terminal region of ArkA and resulting in a more compact conformation. In the final domain-peptide complex, ArkA adopts a PPII helix with an extended conformation, and therefore the *cis* proline conformations in the unbound ArkA ensemble would be unlikely to bind to the SH3 partner with high affinity.

Consequently, the kinetics of binding could be affected by the fraction of the unbound peptide ensemble that contains prolines in *cis*. In a previous study, we characterized the binding pathway of ArkA to the Abp1p SH3 domain with all ArkA prolines trapped in the *trans* conformation and identified an intermediate encounter complex ensemble that forms quickly and is stabilized by nonspecific electrostatic and hydrophobic interactions (Gerlach et al., 2020). While a *cis* conformation of ArkA would not be able to form the canonical fully bound state, it is possible that a *cis*-containing peptide might still interact with the SH3 domain nonspecifically in an encounter complex. It is unknown whether an encounter complex formed with a *cis* conformation of ArkA might be less stable or shorter lived than the *trans* encounter complex, and this is something we hope to investigate in future work. Because many IDP binding sequences, including most SH3 binding regions, contain multiple proline residues, the interactions involving *cis* conformations need to be considered in order to fully understand the complete binding pathway for these proteins.

## 5 Data Availability Statement

The raw data supporting the conclusions of this article will be made available by the authors upon request.

## 6 Author Contributions

KAB and RS designed the simulations. JA, KH, AC, and RE ran and analyzed the simulations. KAB and JA designed the analysis. JA produced the figures. VJM performed and analyzed the CD experiments. EJS designed the CD experiments. KAB and JA wrote the manuscript. All authors revised and approved the final version of the manuscript.

## 9 Acknowledgements

The authors thank Michael Donnelly for computational support. K.A.B. thanks the MERCURY Consortium for mentoring support. The authors thank Alex Holehouse and Paul Nerenberg for helpful discussions.

## 10 Supplementary Material

Included as a separate pdf.

