## Supplemental Material for "Gaussian accelerated molecular dynamics reveals that a proline-rich signaling peptide frequently samples *cis* conformations when unbound"

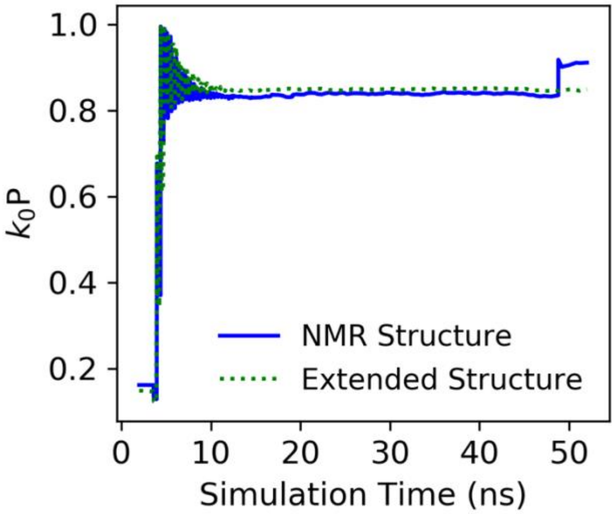


**Figure S1.** Total potential acceleration constant ($k_0$P) from ArkA GaMD equilibrium pre-production simulations (52 ns, 2 fs step). The $k_0$P value determines how "shallow" a given conformational minima becomes if its potential energy is below the defined threshold required to receive an energy boost from GaMD.

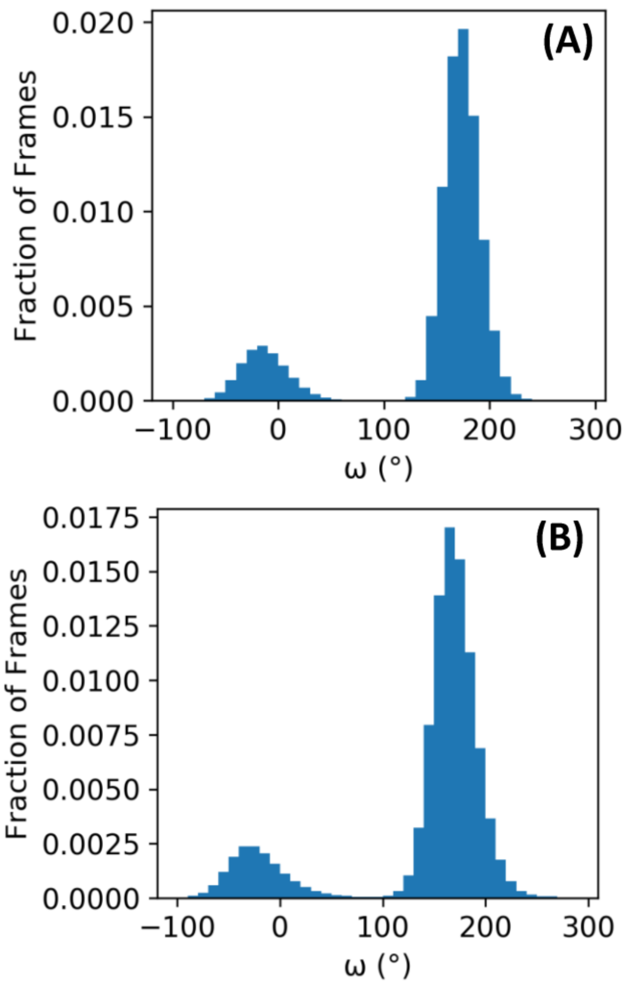


**Figure S2.** Omega angle distributions for the first proline in the KPTP peptide from two independent simulations using the default **(A)** and lowered **(B)** Amber forcefield peptide bond energy barriers (respectively).

**Table S1.** Average proline *cis* sampling percentages for KPTP and ArkA in the GaMD simulations.

| Proline | % in *Cis* in KPTP: default barrier | % in Cis in KPTP: lowered barrier | % in *Cis*: extended structure simulations | % in *Cis*: NMR structure simulations |
|---|---|---|---|---|
| P(2) | 14 ± 7 | 14 ± 1 | 19 ± 0.5 | 14.8 ± 0.6 |
| P(0) | 10 ± 7 | 9.8 ± 0.5 | 17 ± 1 | 13 ± 2 |
| P(-1) | - | - | 14 ± 1 | 10 ± 2 |
| P(-2) | - | - | 12.8 ± 0.7 | 8 ± 2 |
| P(-4) | - | - | 16.5 ± 0.9 | 13 ± 1 |

**Table S2.** Average proline isomerization frequencies between *cis* and *trans* for both KPTP and ArkA systems.

| Proline | KPTP default barrier (flips/ns) | KPTP lowered barrier (flips/ns) | ArkA extended structure (flips/ns) | ArkA NMR structure (flips/ns) |
|---|---|---|---|---|
| P(2) | 0.13 ± 0.03 | 2.4 ± 0.1 | 27.3 ± 0.3 | 11.1 ± 0.4 |
| P(0) | 0.18 ± 0.07 | 3.9 ± 0.2 | 18.7 ± 0.2 | 7.3 ± 0.5 |
| P(-1) | - | - | 22.6 ± 0.5 | 10.2 ± 0.8 |
| P(-2) | - | - | 37 ± 2 | 15 ± 3 |
| P(-4) | - | - | 26.7 ± 0.5 | 9.1 ± 0.3 |

**Table S3.** Average proline ring puckering states percentages in the GaMD ArkA ensemble

| Proline | % in up pucker | % in down pucker | % in planar |
|---|---|---|---|
| P(2) | 36.3 ± 0.4 | 40.8 ± 0.3 | 22.9 ± 0.6 |
| P(0) | 38.0 ± 0.3 | 38.3 ± 0.4 | 23.7 ± 0.6 |
| P(-1) | 36.2 ± 0.4 | 39.8 ± 0.3 | 24.0 ± 0.6 |
| P(-2) | 35.0 ± 0.3 | 41.9 ± 0.5 | 23.1 ± 0.7 |
| P(-4) | 38.1 ± 0.8 | 39 ± 1 | 23.2 ± 0.7 |

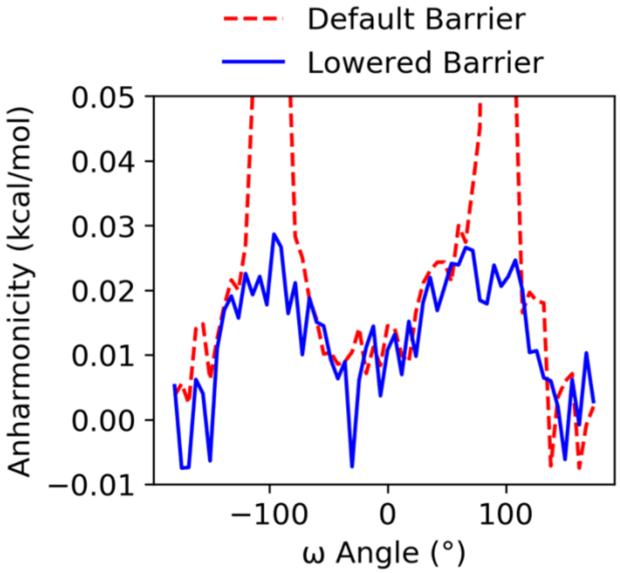


**Figure S3.** GaMD anharmonicity values for first proline omega angle in KPTP default (red dashed line) and lowered barrier (blue solid line) simulations.

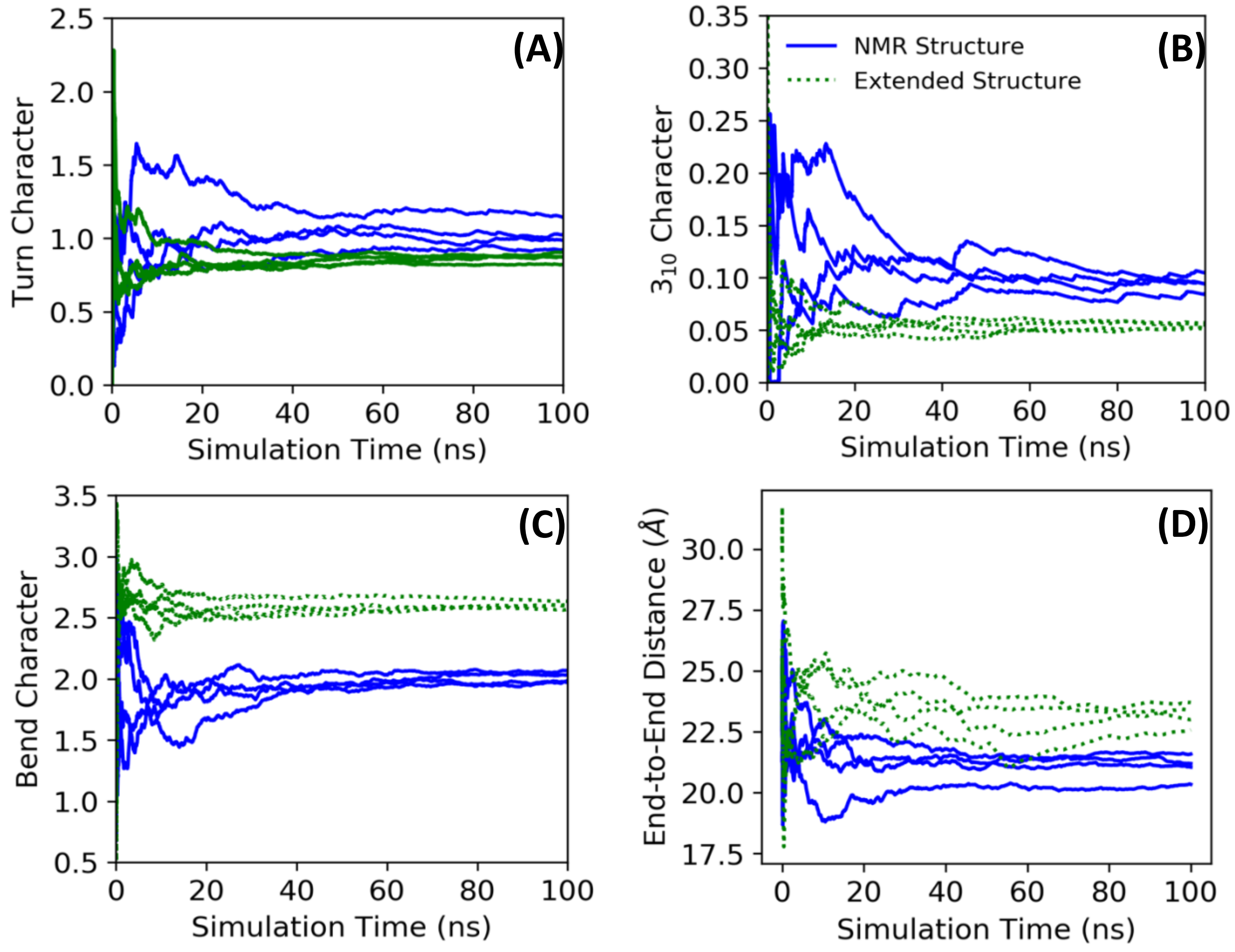


**Figure S4.** Turn percentage **(A)**, $3_{10}$ helix percentage **(B)**, bend percentage **(C)**, and end-to-end distance **(D)** running averages for the ArkA extended (green dotted lines) and NMR (blue solid lines) starting structure GaMD simulations. Running averages were calculated using every $100^{th}$ frame (10 ps) of the one microsecond simulations. Each line represents an independent simulation.

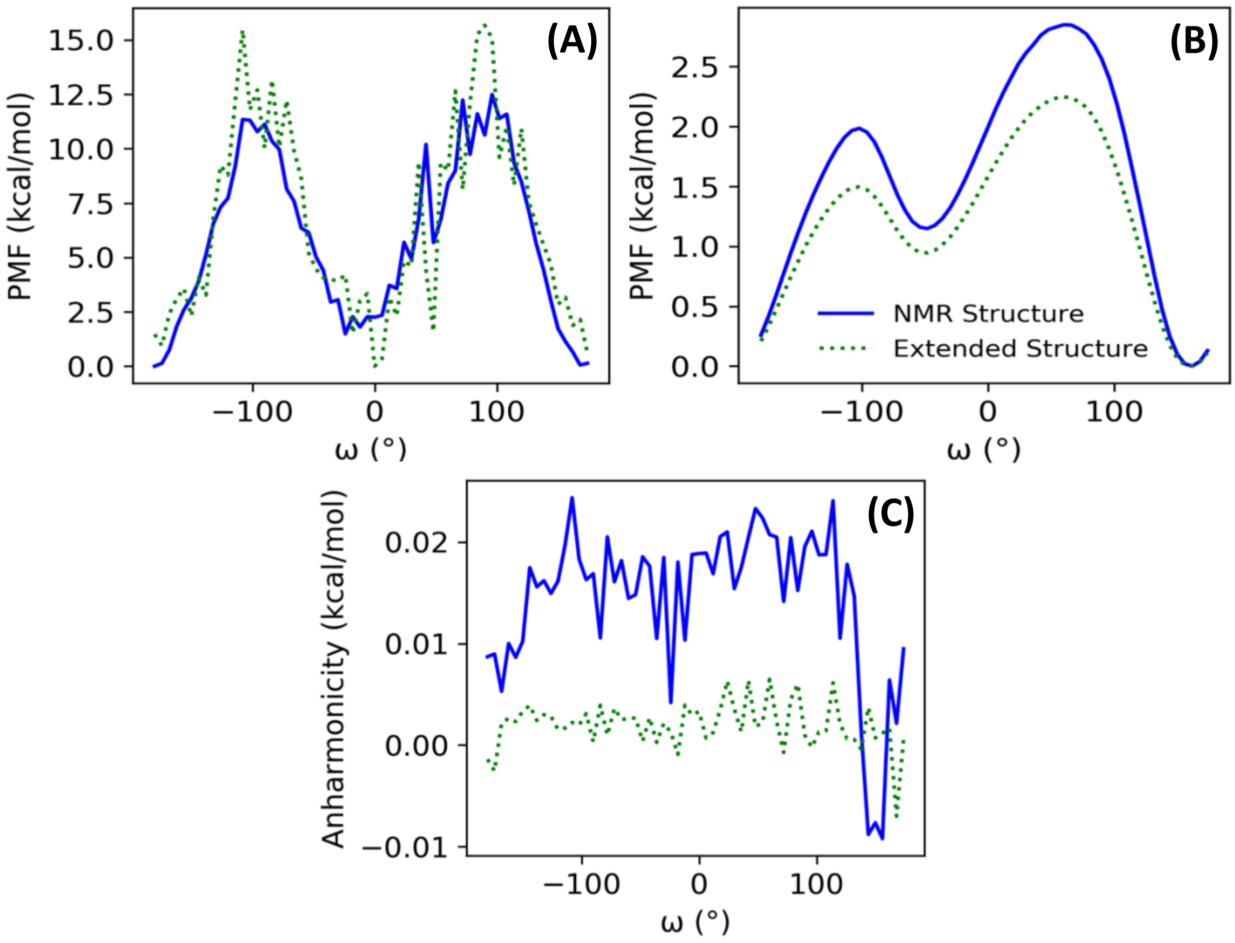


**Figure S5. (A)** PMF free energy landscapes for GaMD ArkA simulations for the first proline reweighted using **(A)** cumulant expansion on the second order and **(B)** Maclaurin series expansion. **(C)** GaMD anharmonicity values for first proline omega angle in GaMD ArkA simulations.

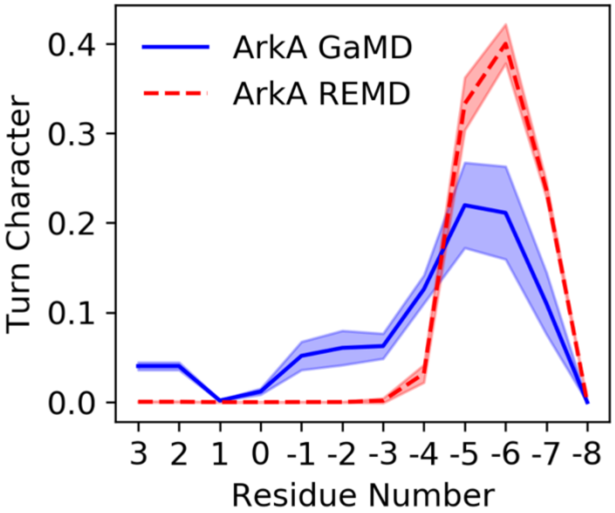


**Figure S6.** Average turn character for the combined NMR and extended structure ArkA GaMD and REMD (Gerlach et. al., 2020) simulations. The shaded regions represent the standard deviations between the respective independent simulations.

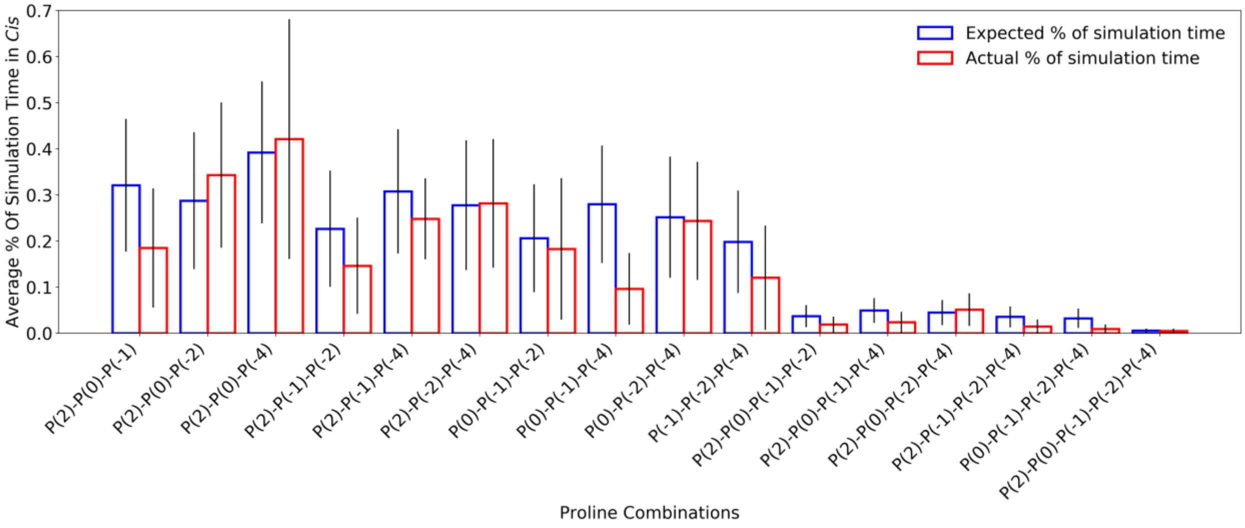


**Figure S7.** Proline *cis* state correlations for three or more residues in the ArkA GaMD simulations. Error bars represent the standard deviation between independent simulations. Calculated expected percentages (on the left in blue) are calculated based on the individual *cis* percentages for each proline assuming all proline residues sample *cis* independently of each other.
